# LOXL2 Deletion Triggers TMJ Osteoarthritis While Overexpression Protects Against NF-κβ-Induced Chondrocyte Apoptosis

**DOI:** 10.1101/2025.05.12.653519

**Authors:** Rajnikant Dilip Raut, Chumki Choudhury, Faiza Ali, Amit kumar Chakraborty, Mohammed Moeeduddin Ahmed, Cheyleann Del Valle-Ponce De Leon, Harshal V. Modh, Pushkar Mehra, Yuwei Fan, Alejandro Almarza, Manish V. Bais

**Author notes:** **Corresponding author: Manish V. Bais:**, Associate Professor of Translational Dental Medicine, Boston University Henry M. Goldman School of Dental Medicine. Equally contributed to work (Rajnikant Dilip Raut, Chumki Choudhury).

## Abstract

Temporomandibular joint osteoarthritis (TMJ-OA) affects a significant proportion of the population worldwide. However, there has been no substantial progress in the development of FDA-approved drugs for treatment due to a lack of understanding of the specific factors regulating key TMJ-OA molecular mechanisms. Lysyl Oxidase Like-2 (LOXL2) promotes knee joint cartilage protection, and it is downregulated in TMJ-OA animal model. We evaluated the role of LOXL2 in TMJ cartilage, its molecular mechanism and gene networks using *in vivo Loxl2* knockout mice (*Acan-Cre; Loxl2*^*flox/flox*^) and *ex vivo* goat TMJ cartilage. Our results show that *Loxl2* knockout in mice cartilage upregulates *Il1b, Mmp9, Mmp13, Adamts4*, and *Adamts5*, whereas it reduces the levels of aggrecan and proteoglycan. *Loxl2* deleted TMJ cartilage show a higher enrichment of inflammatory response, TNFA signaling via NF-kB, extracellular matrix (ECM), and collagen degradation pathway network. Conversely, LOXL2 treatment reduces interleukin-1 beta (IL-1β)-induced expression of *Mmp13*, protects mitochondrial function and ECM from degeneration. Importantly, LOXL2 attenuates IL-1β-induced chondrocyte apoptosis via phosphorylation of NF-κB and expression of pain-related gene *PTGS2* (encodes COX2). Taken together, *Loxl2* knockout mice exacerbate TMJ-OA through cartilage/ECM degradation, mitochondrial dysfunction, chondrocyte apoptosis, and inflammatory gene expression, whereas LOXL2 treatment mitigates these effects.

**Graphical Abstract:** 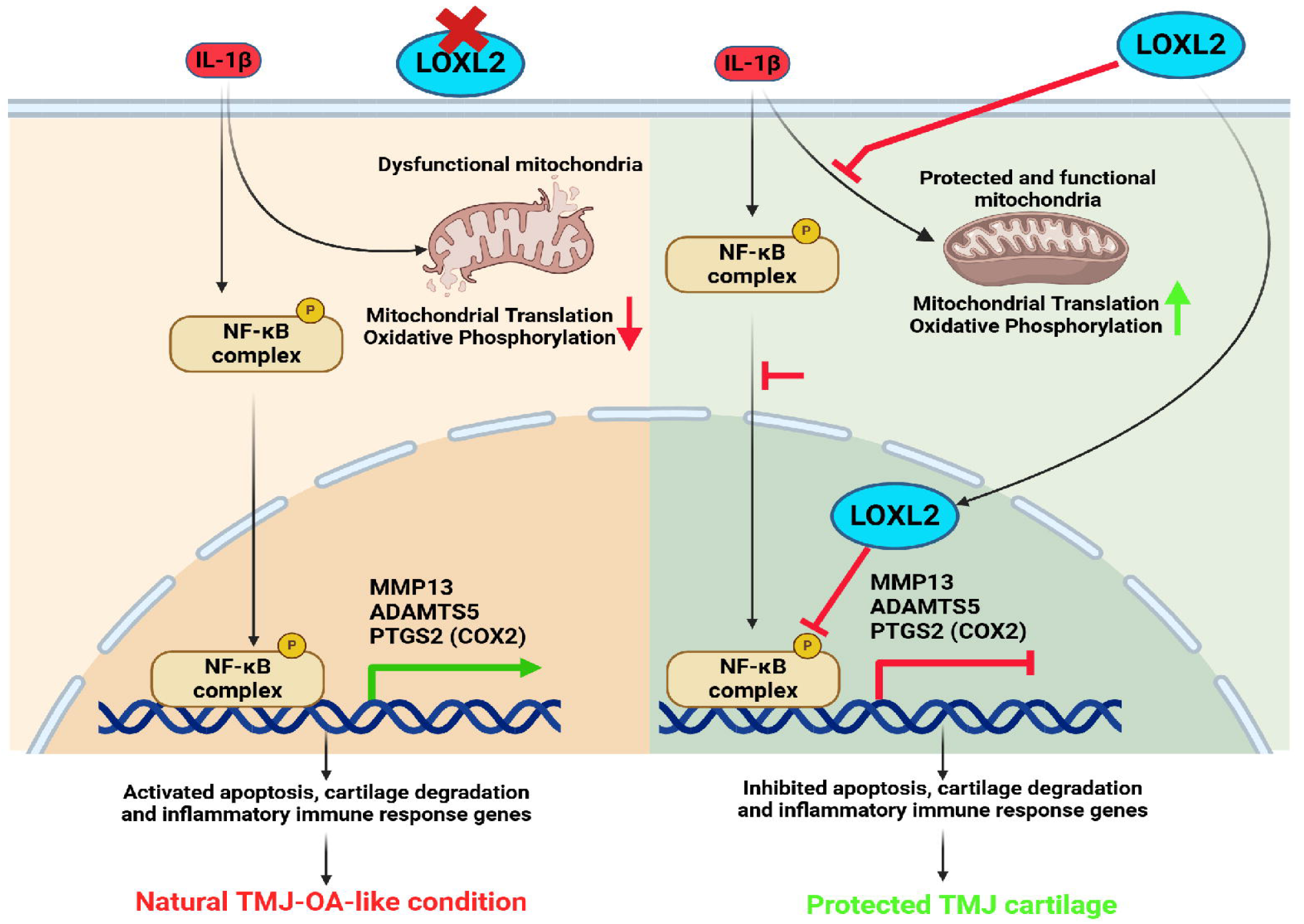

## Introduction

Temporomandibular disorders (TMDs) are prevalent and debilitating conditions that affect 5-12% of the US population. In the United States, TMDs are associated with an annual cost of $4 billion ^1,2^. Temporomandibular joint osteoarthritis (TMJ-OA) is characterized by degenerative cartilage that can cause substantial structural and functional changes. Damage to this cartilage can initiate biochemical changes and structural modifications in the TMJ condyle, promoting TMJ-OA that may not be treatable by maxillofacial surgery. TMJ condylar cartilage has a limited capacity for regeneration. Specific molecular cues promote mesenchymal stem cells or chondroprogenitors to regenerate the TMJ and knee joint articular cartilage ^3-7^. The TMJ is a unique joint, as it has both superficial fibrocartilage and mandibular middle condylar cartilage, compared to the knee and other cartilages. Moreover, TMJ has distinct mechanisms of OA progression compared to other joints ^8-11^. Although the TMJ is highly susceptible to OA and pain, no FDA-approved agents promote TMJ fibrocartilage protection or regeneration due to a lack of mechanistic understanding. Thus, the identification of disease-modifying candidates could be a breakthrough therapy.

Our earlier studies identified, for the first time, that lysyl oxidase like-2 (LOXL2) could prevent cartilage damage and promote an anabolic response ^12-14^. LOXL2 is an extracellular enzyme that catalyzes the oxidative deamination of peptidyl lysine residues, thereby enhancing the synthesis of lysyl-derived collagen and elastin crosslinks in the extracellular matrix (ECM). It helps to maintain the tensile strength and structural integrity of numerous tissues. In addition, adenovirus-delivered LOXL2 in transgenic mice showed an anabolic effect on mouse knee joint articular cartilage at both the molecular and functional levels ^14^. LOXL2 upregulates genes encoding anabolic proteins such as Collagen Type II Alpha 1 chain (COL2A1), sex-determining region Y-box 9 (SOX9), and aggrecan (ACAN), as well as the epigenetic regulator Lysine Demethylase 6B (KDM6B) in human chondrocytes *in vitro*. Furthermore, adenoviral delivery of LOXL2 promotes an anabolic response in the TMJ cartilage of chondrodysplasia (Cho/+) mice, a model of progressive TMJ degeneration. In addition, other groups have shown that LOXL2 promotes the biomechanical properties of cartilage^15,16^. However, if endogenous LOXL2 plays an essential role in preserving TMJ cartilage, its mechanisms in the unique TMJ joint and potential future TMJ-OA translational therapies are unknown.

Here, the importance of LOXL2 was studied by deleting endogenous *Loxl2*, followed by global RNA-seq and histological analyses. Next, LOXL2 overexpression was investigated in murine TMJ cartilage explants, which showed that LOXL2 reverses IL-1β-induced inflammatory changes. The goat TMJ is similar to the human TMJ and is a novel model for potential TMJ regenerative therapeutics ^17,18^. Considering the translational significance of LOXL2 in TMJ regenerative medicine, the role of LOXL2 was investigated in a goat TMJ ex vivo model, which showed that LOXL2 attenuated various processes, including IL-1β induced MMP13, mitochondrial dysfunction, and NF-κβ phosphorylation, which promotes chondrocyte apoptosis.

## Materials and Methods

### MJ Cartilage-specific LOXL2 knockout mice

The *Loxl2* floxed (fl) mice were obtained from Dr. Cano’s laboratory (Madrid, Spain) ^19^ and crossed with aggrecan promoter specific ERT2 inducible Cre-recombinase Acan^tm(IRES-CreERT2)^ or the Acan-Cre^ERT2^ mouse line (Jax #019148) to generate Acan-Cre;*Loxl2*^flox/flox^. Genotyping was performed with specific primers using polymerase chain reaction (PCR) ^19^ and Real time quantitative PCR (RT-qPCR) (Transnetyx Inc.). For tamoxifen-induced deletion of *Loxl2*, one 100-μL intraperitoneal Tamoxifen (75 mg/kg in corn oil) was administered daily for five consecutive days, followed by a maintenance dose of a single injection every month. Six-month-old Acan-Cre;*Loxl2*^fl/fl^ mice were divided into two groups and intraperitoneally injected with either vehicle or tamoxifen (n=8 vehicle, four males and four females) (n=12 tamoxifen, six males and six females). The mice were sacrificed after four months, followed by TMJ histology and RNA sequencing.

### Histology and immunostaining

TMJ joints from *Loxl2* transgenic mice were fixed in 4% paraformaldehyde for 24 hours, decalcified in 10% EDTA (pH 7.4) for 21 days, fixed in paraffin, and subjected to histological analysis and immunostaining. Staining with Safranin-O/Fast Green (American Mastertek Inc., Lodi, CA, USA) was performed as described ^13^. Three slices from four mice per group were deparaffinized, immunostained with specific antibodies to identify LOXL2, ACAN, and MMP13 (Abcam), and visualized with HRP-linked anti-rabbit antibodies. The stained tissues were scanned using a digital slide scanner (Panoramic MIDI; 3D Histech, Budapest, Hungary).

### Ex-vivo goat TMJ cartilage cell culture and adenoviral LOXL2 transduction

Fresh goat TMJ were obtained from local abattoirs (6-9 months of age, female). As per the protocol developed by the Almarza lab, the condylar fibrocartilage was digested with 200U/mL collagenase type II for 1 hour, and the superficial layer cartilage (SLC) was separated from the medial layer cartilage (MLC) of the fibrocartilage. After one hour of digestion, the SLC were peeled, minced, and digested in 200 U/mL collagenase type II (Worthington) for another 3 hours. The CL was separated from the TMJ bone using a scalpel blade and digested in 200 U/mL collagenase type II (Worthington) at 37°C and 5% CO2 with mechanical agitation (rocker, approximately 0.4 Hz). Superficial layer-derived cells (SLC) and middle cartilage layer-derived cells (MLC) were then plated in a culture medium for another 24 hours. Finally, the cells were frozen in dimethyl sulfoxide (DMSO) and shipped to Bosbreaton University. SLC and MLC were cultured in DMEM supplemented with 10% FBS under standard conditions (37°C, 5% CO2). Cells were used at passages 3-5. Treatments included vehicle, IL-1β (10ng), Ad5-LOXL2 (5.2×108 PFU/ml), and a combination of Ad5-LOXL2 and IL-1β. IL-1β treatment lasted for one hour, whereas Ad5-LOXL2 was incubated overnight with the cells for transduction. In the combined treatment group (Ad5-LOXL2+IL-1β), cells were transduced overnight and treated with IL-1β for one hour the following day. After treatment, cells were harvested for RNA sequencing and downstream analysis.

### RNA-sequencing and bioinformatics analyses

Total RNA was extracted from all samples using the TRIzol protocol according to the manufacturer’s instructions (Qiagen). The extracted RNA samples were subjected to RNA sequencing using Novogene (Sacramento, California, USA). All the samples were tested for quality before library construction. Only samples that passed the quality check were selected based on the RNA Integrity Number (RIN). Paired-end RNA sequencing was performed using the Illumina high-throughput sequencing platform. Raw FASTQ sequencing reads were assessed for quality control and trimmed or preprocessed using an in-house Novogene Perl script. The filtered data were mapped against the reference genomes of Mus musculus (mm10) and Capra hircus (ncbi_capra_hircus_gcf_001704415_2_ars1_2). Feature Counts (v1.5.0-p3) were used to quantify the number of raw reads that mapped to each gene. Differential gene expression (DGE) analysis was performed using the DESeq2 package in the R/Bioconductor software. Functional enrichment analysis was performed using the gene set enrichment analysis (GSEA-v4.3.2) software to evaluate the biological processes or hallmark pathways implicated. Heatmaps were generated using the ComplexHeatmap package in R. Volcano plots for differentially expressed genes were generated using the ggplot2 package in R. Box/bar plots were generated using GraphPad PRISM software (v10.1.0). Ingenuity pathway analysis (IPA) was performed using IPA software (Qiagen).

### Confocal Microscopy for chondrocyte apoptosis, mitochondrial dynamics, and mitophagy

To investigate the effects of different treatments on mitochondrial dynamics, goat cells were exposed to the vehicle, IL-1β, Ad5-LOXL2, or a combination of Ad5-LOXL2 and IL-1β. MitoTracker Red-CMXros (Thermo Fisher Scientific) dye was used to stain mitochondria. DAPI was used to stain the nuclei, allowing for detailed visualization of mitochondrial distribution using confocal microscopy at 63x magnification. To investigate the effect on mitophagy, the cells were treated with a p62-mediated mitophagy inducer (MedChemExpress # HY-115576) (1uM) alone or in combination with Ad5-LOXL2. We also stained the cells with DAPI and FITC-annexin V in combination with PE-anti phospho NF-κβ (BD #558423) to observe the apoptotic effect induced by IL-1β and the protective role of LOXL2. Cell fixation/permeabilization and intercellular staining were performed using a True-Nuclear Transcription Factor Buffer Set (BioLegend).

### Flow cytometry and apoptosis assay

Goat TMJ cells were grown in a monolayer and infected with Ad5-Empty or LOXL2 (5.2×10^8^PFU/ml) in respective groups overnight, followed by stimulation with IL-1β for one h. FITC-annexin V assay was performed according to the manufacturer’s instructions to determine the percentage of chondrocytes undergoing apoptosis. After treatment, TMJ cells were collected, washed twice with cold PBS, and resuspended in 100□μL of binding buffer. FITC annexin V (Thermo # A13199) or anti-MMP13 antibodies (5 μL per tube) were added in the respective groups and incubated for 15 min at 25°C in the dark. Intercellular staining with PE anti-phospho NF-κB (p65) (BD #558423) was performed using the True-Nuclear™ Transcription Factor Buffer Set (BioLegend #424401) following the manufacturer’s protocol and examined using a five-laser 64-color Cytek Aurora spectral flow cytometer.

## Results

### Aggrecan promoter-specific Loxl2 knockout promotes TMJ-OA in mice

To understand the role of LOXL2 during TMJ-OA, we reanalyzed the available RNA sequencing data ^20^ and detected that LOXL2 was significantly downregulated in the rabbit TMJ-OA model compared to the healthy TMJ, whereas Il1b (IL-1β) expression was significantly increased (Fig. 1a). This shows that LOXL2 and IL-1β could have an inverse relationship in TMJ-OA. Increased expression of IL-1β is a key event in the progression of natural OA ^21^. To evaluate whether LOXL2 loss-of-function promotes progressive degenerative changes in TMJ condylar cartilage, aggrecan promoter-specific tamoxifen-inducible *Loxl2* knockout mice were generated (Fig. 1b). *Loxl2* knockout mice showed a reduction in LOXL2, ACAN, and proteoglycans and an increase in MMP13 (Fig. 1c).

**Fig. 1:**
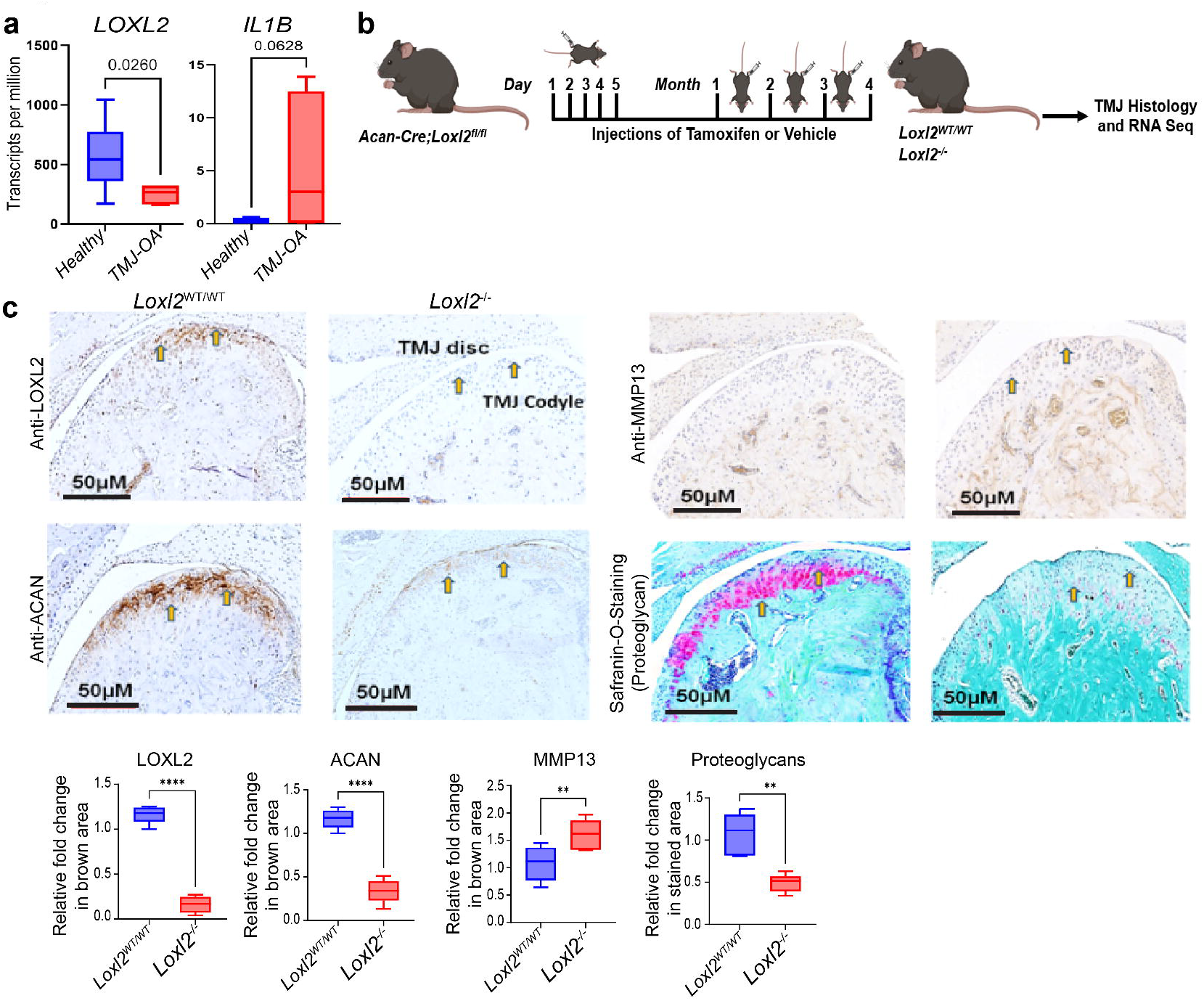
Cartilage-specific Loxl2 knockout in mice promotes TMJ-OA. **a)** LOXL2 expression in rabbits’ healthy TMJ and TMJ-OA analyzed from available data (p-value calculated using Wilcoxon rank sum test). b) Experimental design for tamoxifen-induced *Loxl2* knockout mice; *Acan-Cre;Loxl2*^*fl/fl*^ mice were injected with vehicle or tamoxifen daily for five consecutive days, followed by one injection per month for 4 months to generate *Loxl2*^*WT/WT*^ or *Loxl2*^*-/-*^, respectively. **c)** Immunohistochemistry shows reduced levels of LOXL2 and ACAN, whereas increased levels of MMP13 in the *Loxl2* knockout mice; Safranin-O staining shows decreased levels of proteoglycans in the *Loxl2* knockout mice. *p<0.05, **p<0.01, ***p<0.001, ****p<0.0001.

### Aggrecan promoter-specific Loxl2 knockout promotes TMJ-OA related molecular changes in mice

RNA sequencing (RNA-Seq) and differential gene expression (DGE) analysis identified 4242 dysregulated genes (padj<0.05) in *Loxl2* deleted mice, of which 2240 protein-coding genes were upregulated and 2002 were downregulated (Fig. 2a and Table S1). Interestingly, *Loxl2* deletion enhanced the expression of Il1b (IL-1β), *Mmp9, Mmp13, Adamts4*, and *Adamts5*, which are widely known for their roles in OA (Fig. 2b). GSEA showed higher enrichment for inflammatory responses and TNF-α signaling via NF-κβ, extracellular matrix degradation, and collagen degradation networks (Fig. 2c). The *Loxl2* deletion TMJ cartilage signature had a higher enrichment of genes related to mast cell activation, macrophage activation, neuroinflammatory response, cytokine production, and IL6 production, contributing to OA and pain (Fig. 2d). The *Loxl2* deletion also inhibited specific genes and pathways related to mitochoria, affecting mitochondrial translation and oxidative phosphorylation (Fig. 2e). Ingenuity pathway analysis (IPA) predicted that *Loxl2* knockout promoted cartilage degradation, cleavage of collagen and proteoglycan, chondrocyte death, and activation of the IL-1β network (Fig. S1a, b).

**Fig. 2:**
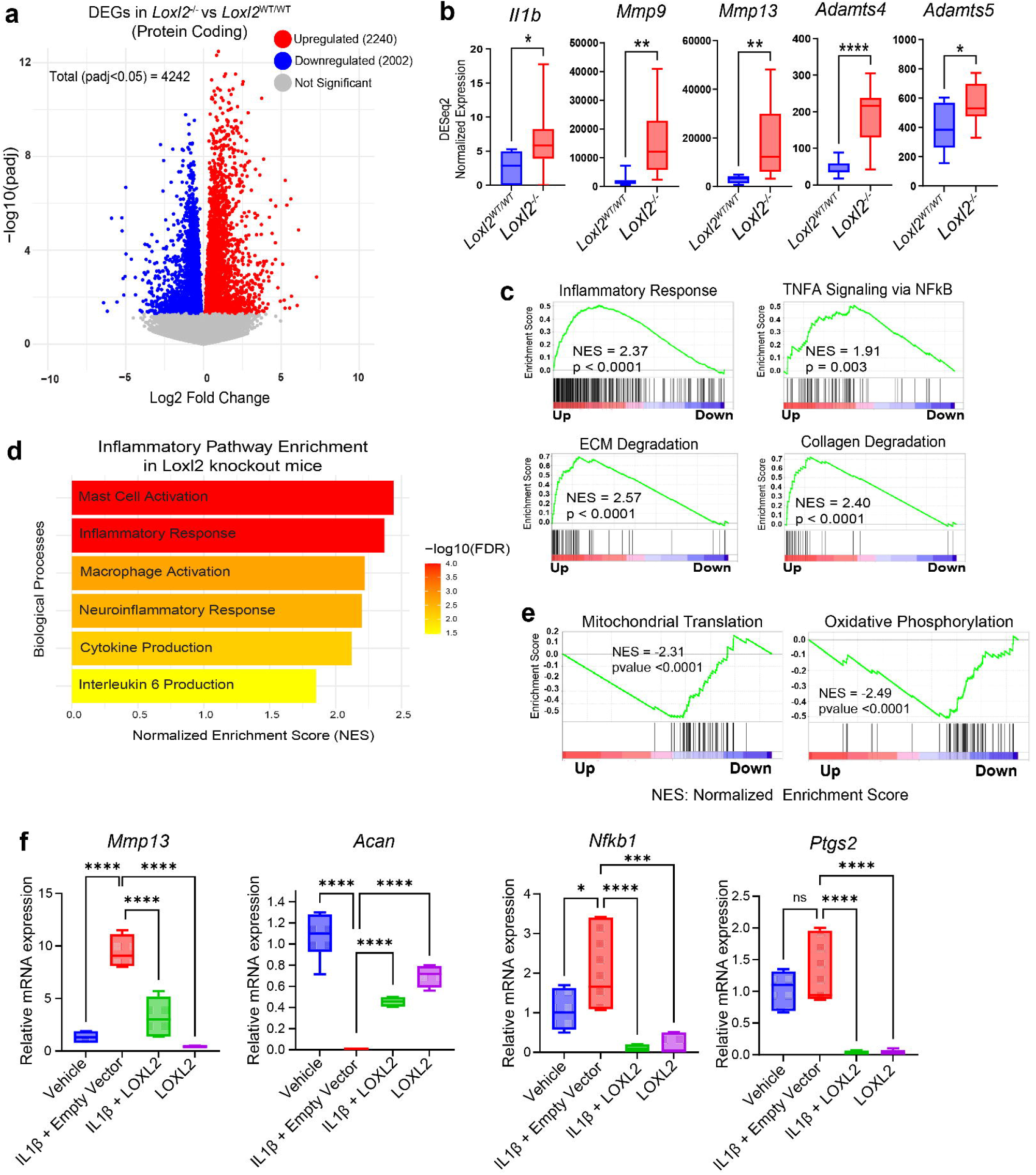
RNAseq analysis identified *Loxl2* knockout shows TMJ-OA like molecular signature. **a)** Volcano plot showing total differentially expressed genes with *Loxl2* knockout in mice; red and blue dots represent significantly upregulated and downregulated genes, respectively, whereas grey dots represent genes with |adjusted p-value>0.05|. **b)** Boxplots representing elevated expression *Il1b* (IL-1β), *Mmp9, Mmp13, Adamts4*, and *Adamts5* in the RNA-seq data of *Loxl2* knockout mice (p-values were calculated using default Wald test by DESeq2 algorithm). cC GSEA plots showing increased enrichment of inflammatory response and ECM/Collagen degradation gene sets with *Loxl2* knockout. **d)** Increased enrichment of inflammatory pathway gene sets in *Loxl2* knockout mice. **e)** GSEA analysis shows *Loxl2* knockout mice inhibits mitochondrial translation (MT) and oxidative phosphorylation (OP). NES: Normalized enrichment score. **f)** RT-qPCR analysis of mouse TMJ explants treated with vehicle, IL-1β, IL-1β+LOXL2, and LOXL2 showed affected mRNA expression of *Mmp13, Nfkb1, Ptgs2*, and *Acan* (p-values were calculated using one-way ANOVA). p<0.05, **p<0.01, ***p<0.001, ****p<0.0001.

### Adenoviral LOXL2 treatment in mice TMJ explants reverse OA-specific molecular signatures

Increased expression of IL-1β is a key event in the progression of natural OA ^21^. Next, to evaluate whether LOXL2 could reverse IL-1β-induced OA-like changes, fresh murine TMJ cartilage explants were treated with vehicle, IL-1β, LOXL2, or a combination of IL-1β and LOXL2 in respective groups. IL-1β treatment promoted *Mmp13, Nfkb1*, and *Ptgs2*, which was significantly reduced by LOXL2 treatment. In contrast, IL-1β reduced *Acan*, which was restored by LOXL2 treatment (Fig. 2f). Hence, we concluded that the loss of *Loxl2* promotes inflammation, leading to natural OA-like changes and pain in the mouse TMJ, which could be rescued by LOXL2 treatment.

### Reduction in IL-1β induced degenerative effects in goat TMJ cartilage by LOXL2 overexpression

The TMJ varies in structure and function compared to other joints and has two layers of cartilage including superficial layer cartilage (SLC) and medial layer cartilage (MLC)^22^: SLC and MLC are distinct which has been shown in previous studies. Therefore, we independently evaluated the effect of LOXL2 treatment on OA induced by IL-1β in goat TMJ cartilage explant overall (Fig. 3a) and in both SLC and MLC (Fig. 3b).

**Fig. 3:**
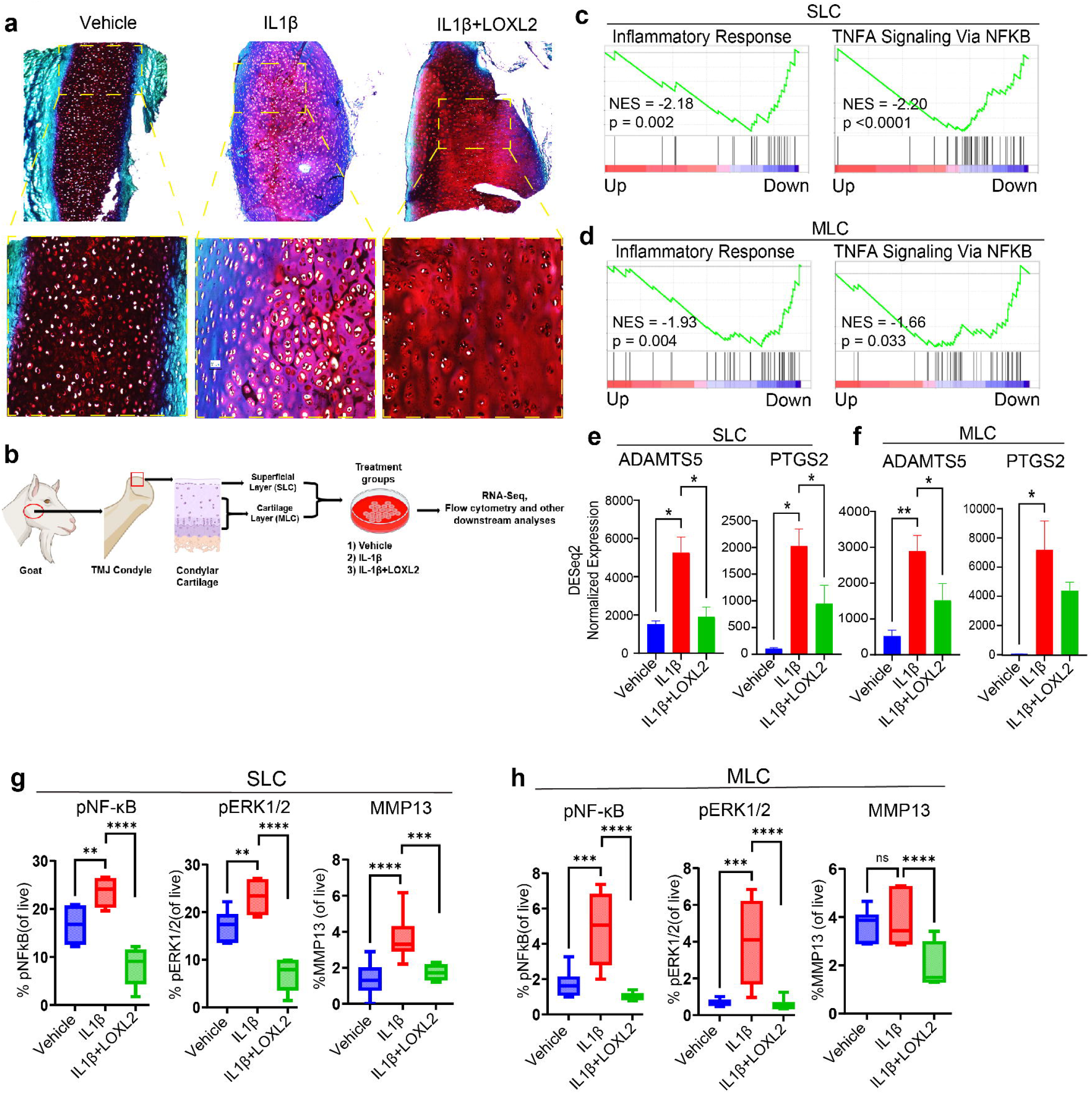
LOXL2 treatment in the goat TMJ cartilage protects from OA. **a)** Histology analysis showed increased proteoglycans in the LOXL2-treated group. b) Experimental design of *ex vivo* goat TMJ cartilage cells isolation and culture, LOXL2 treatment, and downstream analyses. **c-d)** RNAseq/DGE analysis followed by GSEA shows a significant reduction in inflammatory response and TNF alpha signaling via NF-κβ gene sets in both SLC and MLC after LOXL2 treatment. **e-f)** Bar plots showing increased mRNA expression of *ADAMTS5* and *PTGS2* with IL-1β treatment, which was attenuated in LOXL2 treated group in SLC and MLC (p-values were calculated using default Wald test by DESeq2 algorithm). **g-h)** Flow cytometry showing MMP13, phospho-NF-κβ, and phospho-ERK1/2 expression. The percentage of MMP13, pNF-κβ, and pERK1 was significantly higher in the IL-1β treated group, which was rescued by LOXL2 treatment (p-values were calculated using a one-way ANOVA test). These results advocate a protective role of LOXL2 in the TMJ-OA. ns>0.05 *p<0.05, **p<0.01, ***p<0.001, ****p<0.0001.

To analyze the overall effect on TMJ cartilage, freshly isolated healthy goat TMJ explants were treated with vehicle, IL-1β, or IL-1β+LOXL2 and subjected to Safranin-O staining. Consistent with IPA findings, IL-1β affected proteoglycan (GAG; Glycosaminoglycan) levels, whereas LOXL2 rescued the proteoglycan (Fig. 3a). Tables S2 and S3 show the total differentially expressed genes in IL-1β+LOXL2 compared to the IL-1β treated group for both SLC and MLC. LOXL2-treated chondrocytes showed significantly reduced enrichment of genes associated with the inflammatory response and TNF-α signaling via NF-κβ (Fig. 3c, d). Moreover, induced expression of key OA was also observed in IL-1β SLC and MLC (Fig. S2a). IPA predicted that LOXL2 treatment resulted in the inhibition of catabolic processes such as cartilage degradation, proteoglycan cleavage, cleavage of collagen fibers, chondrocyte death, and activation of anabolic glycosaminoglycan processes (Fig. S2b). Additionally, LOXL2 treatment reversed the level of IL-1β-induced *ADAMTS5* and *PTGS2* (COX2) (Fig. 3e, f). ADAMTS5 and COX2 are the major cartilage degradation factors and pain inducers. This data highlights that LOXL2 plays a key role in controlling the expression of inflammatory and pain marker genes in TMJ chondrocytes.

### IL-1β-induced degenerative effects, including NF-κβ activation leading to MMP13 induction, are inhibited by LOXL2 in goat TMJ chondrocytes

The activation of NF-κβ is an early event that occurs during the immune response ^23^, and constitutive NF-κβ activation leads to OA-like changes ^24^. Therefore, we performed flow cytometry to analyze NF-κβ phosphorylation (pNF-κβ) and MMP13 protein expression. Interestingly, pNF-κβ levels were increased after IL-1β treatment and decreased after treatment with LOXL2 (Fig. 3g, h; Fig. S3). Studies suggest that NF-κβ is activated through ERK1/2 signaling, where the ERK1/2 pathway activates the IkB kinase (IKK) complex that phosphorylates IkB, resulting in its degradation and subsequent nuclear translocation of NF-κβ ^25^. As expected, phosphorylated ERK1/2 levels were significantly elevated by IL-1β treatment and were restored to normal levels by LOXL2 treatment (Fig. 3g, h; Fig. S3). Furthermore, MMP13 protein levels were increased in IL-1β-stimulated SLC, whereas LOXL2 treatment resulted in a significant MMP13 reduction in both SLC and MLC (Fig. 3g, h; Fig. S3).

Overall, we conclude that IL-1β treatment causes proteoglycan degradation to promote TMJ-OA by inducing pNF-κβ and MMP13, whereas LOXL2 reverses these effects, resulting in TMJ protection.

### LOXL2 preserves IL-1β-induced mitochondrial dysfunction and p62-mediated mitophagy

To delineate the possible mechanism by which LOXL2 protects TMJ cartilage, RNA-seq data and GSEA detailed analysis revealed that gene sets related to mitochondrial translation and oxidative phosphorylation had higher enrichment in LOXL2 treated group, providing a scientific basis for testing whether LOXL2 protects TMJ cartilage by controlling mitochondrial function (Fig. 4a, b). A subset of candidates from the mitochondrial translation (mitochondrial ribosomal subunits) and oxidative phosphorylation (NADH: Ubiquinone Oxidoreductase (Complex I) subunits) gene sets were negatively correlated, as *Loxl2* knockout resulted in their downregulation, whereas adenoviral LOXL2 treatment enhanced their expression significantly (Fig. S4a-d). These data suggested that LOXL2 treatment enhanced mitochondrial protein synthesis and energy production in TMJ chondrocytes.

**Fig. 4:**
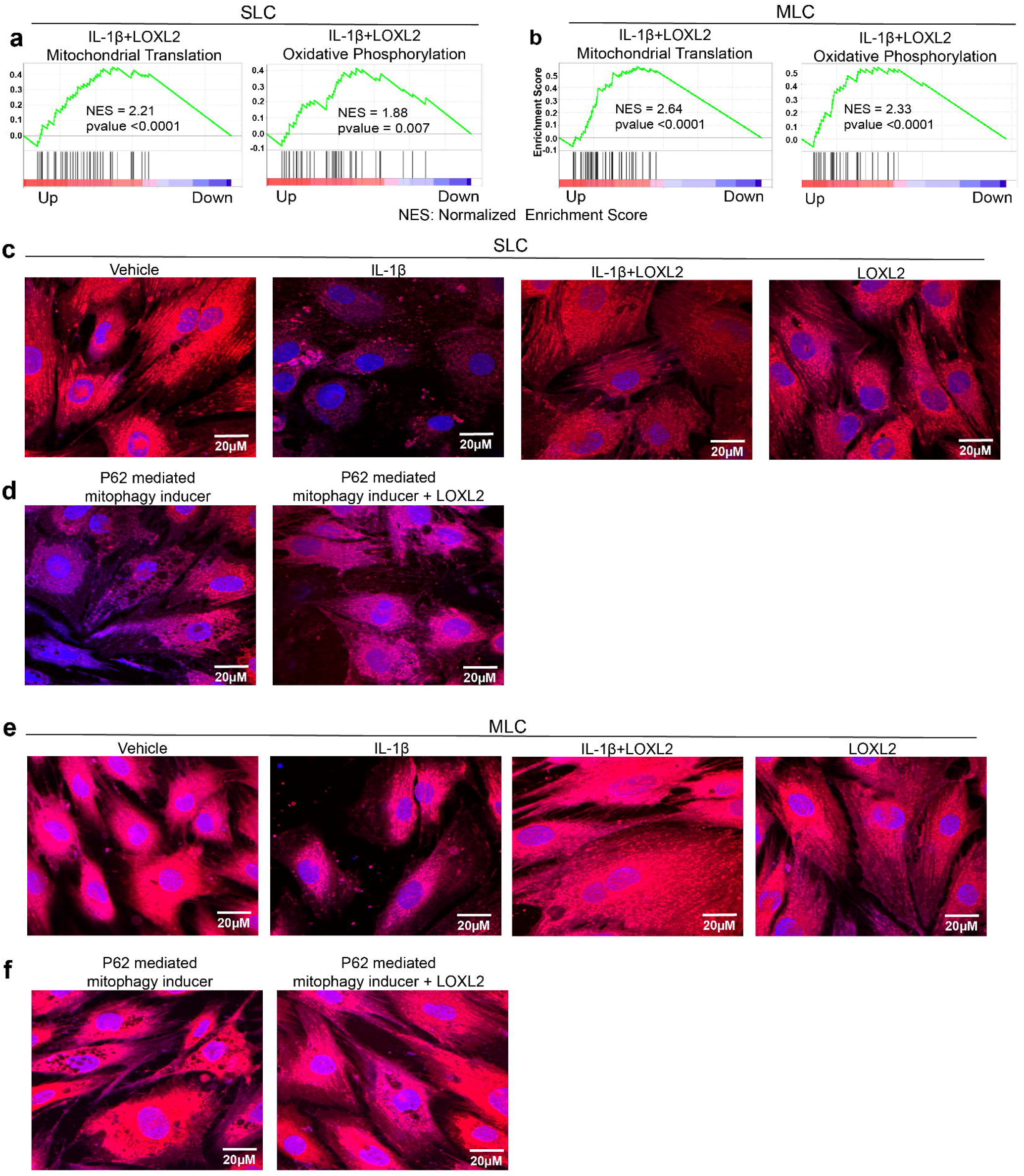
LOXL2 protects and enhances goat TMJ chondrocyte mitochondrial function. **a-b)** GSEA analysis of LOXL2 treated TMJ chondrocytes shows higher enrichment of mitochondrial translation (MT) and oxidative phosphorylation (OP) in SLC and MLC. NES: Normalized enrichment score. **c)** Confocal microscopy showing a reduction in mitochondrial abundance in IL-1β treated SLC, which was preserved in LOXL2 treated group; this effect was similar to the vehicle and LOXL2-only treated groups. **d)** p62-mediated mitophagy inducer showed degenerative phenotype in SLC, which was preserved when treated with LOXL2. **e)** Confocal microscopy showing a reduction in mitochondrial abundance in IL-1β treated MLC, which was preserved in LOXL2 treated group; this effect was similar to the vehicle and LOXL2-only treated groups. **f)** p62-mediated mitophagy inducer showed degenerative phenotype in MLC, which was preserved when treated with LOXL2. MitoTracker™ -Red dye was used to stain the active mitochondria. Images were captured at 63X resolution.

Next, we evaluated the association between mitochondrial function and TMJ-cartilage protection. TMJ chondrocytes were treated with vehicle, IL-1β, IL-1β, LOXL2, or LOXL2. IL-1β treatment reduced the mitochondrial abundance, which was protected by LOXL2 (Fig. 4c, e). The mitochondrial number is controlled by mitophagy; however, excessive or dysregulated mitophagy contributes to chondrocyte death and cartilage degradation ^26^. To test this hypothesis, TMJ chondrocytes were treated with a p62-mediated mitophagy inducer, which resulted in a degenerative cellular phenotype (Fig. 4d, f). This effect was not observed when cells were incubated with LOXL2 (Fig. 4d, f). Overall, we concluded that LOXL2 treatment of TMJ chondrocytes protects against the degenerative effects of IL-1β- and p62-mediated mitophagy-induced cartilage degradation, leading to TMJ-OA.

### LOXL2 prevents IL-1β-induced TMJ chondrocyte apoptosis

Mitochondrial dysfunction can activate cell death pathways while limiting cell survival, leading to chondrocyte apoptosis ^27^. Reduced mitochondrial translation and oxidative phosphorylation may lead to decreased ATP production, eventually affecting chondrocyte function and survival by increasing reactive oxygen species (ROS) production. Oxidative stress caused by ROS accumulation can activate the NF-κβ pathway and may contribute to ECM/cartilage degeneration and inflammatory immune response^28,29^. IPA predicted *that Loxl2 knockout in mice increased IL-1*β*-induced* chondrocyte apoptosis via aberrant NF-κβ signaling (Fig. S4a), whereas LOXL2 treatment reversed this effect (Fig. S4b). To validate whether LOXL2 affected chondrocyte apoptosis, we performed flow cytometry with Annexin V immunostaining. IL-1β treatment increased the expression of the chondrocyte apoptosis marker Annexin V. Conversely, LOXL2 treatment significantly reduced apoptosis compared to IL-1β treatment (Fig. 5a, b), suggesting that LOXL2 effectively mitigated TMJ chondrocyte apoptosis.

**Fig. 5:**
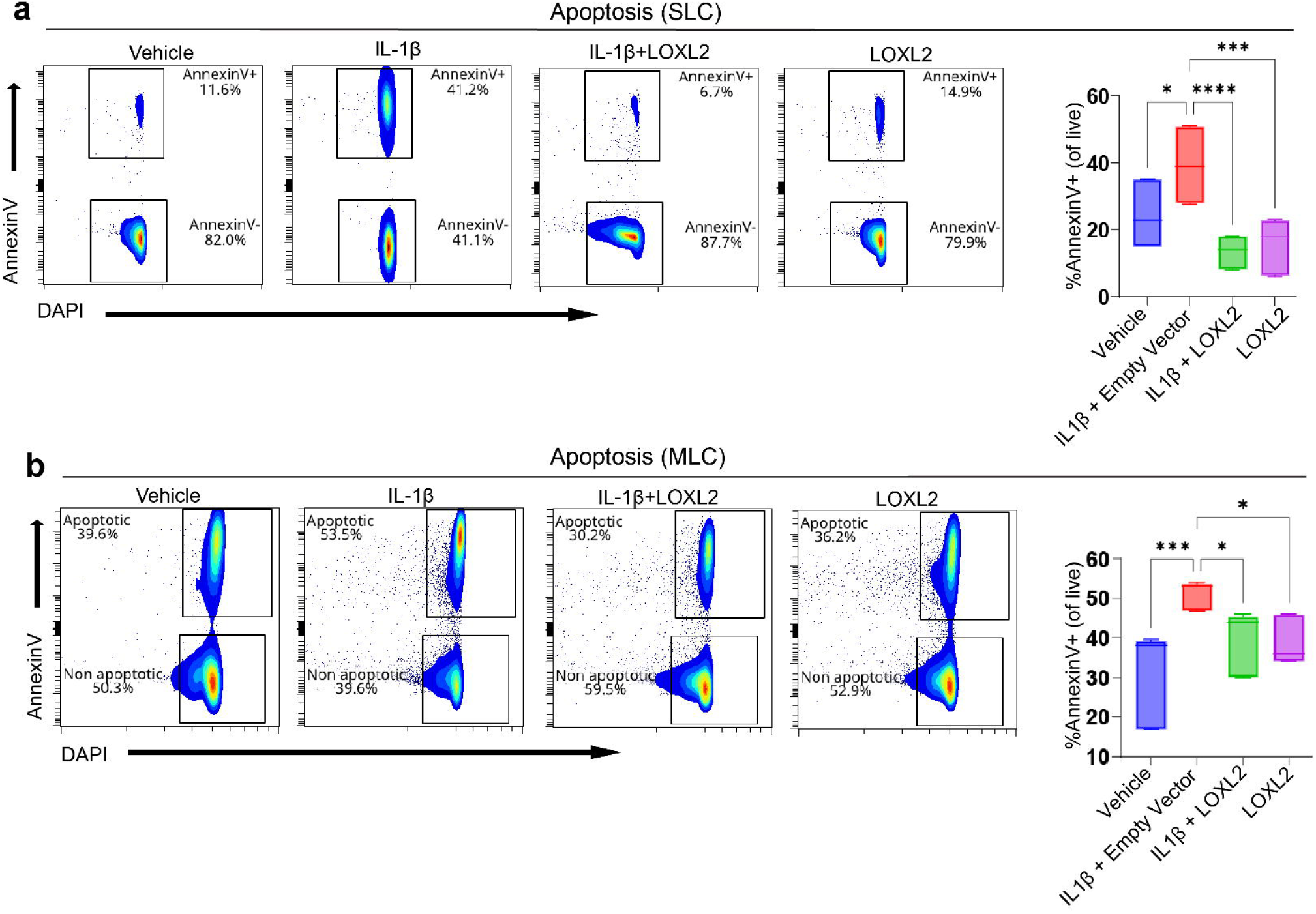
LOXL2 treatment reduces IL-1β-induced chondrocyte apoptosis. **a-b)** Flow cytometry-based detection of apoptotic cell population using AnnexinV immunostaining. IL-1β treatment showed increased apoptosis (%AnnexinV+ cells), while LOXL2 treatment significantly decreased the AnnexinV+ apoptotic cells in both SLC and MLC (p-values were calculated using one-way ANOVA test).

### LOXL2 attenuates IL-1β-phospho-NF-κβ axis mediated apoptosis in goat TMJ chondrotyes

To evaluate the mechanism of protection of chondrocyte apoptosis, we compared IL-1β vs IL-1β, along with LOXL2 treated goat chondroctye transcription signature, which showed that chondrocyte apoptosis was mediated by NF-κβ (Fig. S5a, b). Activation of the NF-κβ pathway leads to degenerative changes and inflammation^28,29^. RelA(p65)/p50 heterodimers is the most common form of NF-κβ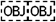. We assessed the levels of phosphorylated NF-κβ (pNF-κβ/p65/RelA) using flow cytometry, as pNF-κβ is required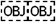 nuclear translocation and transcription of proapoptotic genes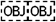. IL-1β treatment elevated pNF-κβ levels, which were reduced in the LOXL2-treated group (Fig. 6a, b). To evaluate the translocation of pNF-κβ in apoptotic chondrocytes, we performed confocal microscopy and visualized the levels of annexin V (green) and pNF-κB (red) (Fig. 6c). We found a significant elevation in Annexin V and pNF-κβ levels after IL-1β treatment, which was inhibited by LOXL2. Hence, we concluded that LOXL2 protects TMJ cartilage from the degenerative effects of IL-1β, chondrocyte apoptosis, and TMJ-OA-like progression.

**Fig. 6:**
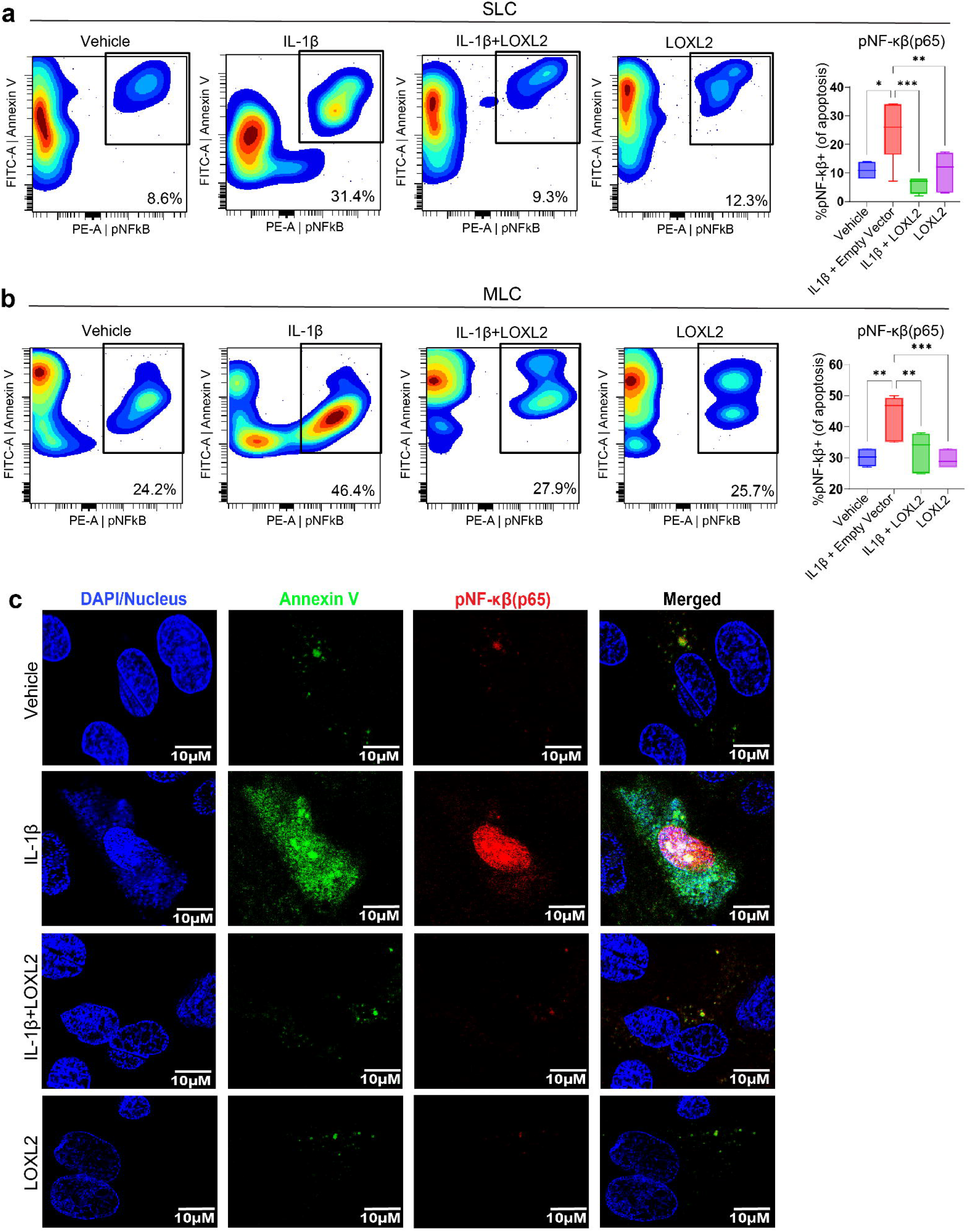
LOXL2 treatment reduces IL-1β-induced chondrocyte apoptosis via inhibiting the NF-κβ pathway. **a-b)** Flow cytometry detects that IL-1β treatment significantly increased the activated pNF-κβ in apoptotic cell population, which was rescued in LOXL2 treatment groups of SLC and MLC (p-values were calculated using a one-way ANOVA test). **c)** Confocal microscopy showing the levels of AnnexinV and pNF-κβ double positive MLC cells were highly increased with IL-1β which were rescued by LOXL2 treatment. Blue represents DAPI-stained nucleus, green represents Annexin V, and red represents activated pNF-κβ translocated to the nucleus. *p<0.05, **p<0.01, ***p<0.001, ****p<0.0001.

## Discussion

The aim of this study was to investigate the mechanistic role of LOXL2 in TMJ-OA. Genetic deletion of LOXL2 in cartilage attenuated aggrecan and proteoglycans while increasing IL-1β, MMP13, and ADAMTS5 levels, leading to progressive TMJ-OA-like changes. LOXL2 treatment protected the TMJ cartilage from the degenerative effects of IL-1β and reduced the inflammatory immune response and TMJ chondrocyte apoptosis. This is the first study to evaluate the variable transcriptomic signatures of IL-1β activation due to loss of LOXL2, whereas LOXL2 overexpression restored specific transcriptional changes, attenuating chondrocyte apoptosis and pNF-κβ.

IL-1β is a pro-inflammatory cytokine contributing to the inflammatory response in the joints ^31,32^ and chondrocyte apoptosis ^27^. IL-1β induces its effects by activating NF-κβ signaling, leading to the transcription of *Mmp13* and *Adamts5* ^33^, leading to cartilage degradation and compromising the structural integrity of cartilage ^34,35^. MMP13 is known for its potent collagenolytic activity, mainly targeting type II collagen, and promoting cartilage degradation and OA ^36^. ADAMTS5 degrades aggrecan and exacerbates OA pathology ^37^. NF-κβ activation worsens disease outcomes in patients with TMJ-OA and discectomy-induced TMJ-OA in mice ^38,39^. NF-κβ is activated through ERK1/2 signaling ^40,41^, and promotes immune response activation ^25^. Moreover, NF-κβ pathway activation induces the expression of ADAMTS5 and aggravates TMJ-OA ^42^. NF-κβ signaling inhibitors have also been developed for OA ^43^.

In addition, *Ptgs2*, a key mediator of PGE2 (prostaglandin E2) production, was also upregulated in *Loxl2* knockout mice and downregulated with LOXL2 treatment. This correlates with previous findings that NF-κβ can also enhance joint injury by inducing COX2, eventually promoting apoptosis of articular chondrocytes, tissue inflammation, and synthesis of catabolic factors ^44^. COX2 is directly associated with synovitis and joint pain in patients with internal derangement or TMJ-OA, rendering its inhibition significant for reducing inflammation and pain in the TMJ ^45,46^. Finally, we validated that LOXL2 treatment restored IL-1β-induced morphological changes, ECM integrity, and apoptosis in chondrocytes, which is critical for understanding the outcome of LOXL2-induced protective effects. Hence, this provides a functional validation that LOXL2 plays a protective role in murine TMJ cartilage, and its loss results in natural TMJ-OA-like conditions.

Looking forward to the clinical translation of LOXL2 as a therapeutic candidate, we analyzed *ex vivo* goat TMJ condylar cartilage explants and cells. As reviewed in our recent publications ^17,18^, goats are ideal large animal models for evaluating TMJ regenerative medicine. One advantage is the accessible joint space, because the zygomatic arch does not hinder access, making its surgical anatomy comparable to that of humans. Another significant advantage is its status as a ruminant species, consuming 12-16 hours daily of chewing food. The successful therapies tested in goats are likely to translate well into humans. Thus, our studies evaluated LOXL2’s potential on IL1β induced chondrocyte apoptosis, mitochondrial dysfunction, pNF-κβ activation and transcription. LOXL2 treatment to goat TMJ chondrocytes resulted in pNF-κβ and pERK1/2 reduction and downregulation of COX2 (*Ptgs2*). Therefore, this strongly supports our hypothesis that LOXL2 protects against inflammatory responses during TMJ-OA.

Furthermore, LOXL2 loss- and gain-of-function gene networks related to mitochondria may be critical for understanding its mechanisms of action. The *Loxl2* knockout resulted in the downregulation of mitochondrial translation and oxidative phosphorylation, whereas these processes were upregulated by LOXL2 treatment. Previously, altered mitochondrial function was shown to contribute to OA development ^27^. Mitophagy and NF-κβ signaling are also initiated as parallel pathways in response to mitochondrial stress^47^. Mitochondrial biogenesis is controlled by p62-mediated mitophagy, which plays a vital role in OA development ^27,48^. Mitophagy activation through PINK1 and Parkin may lead to the clearance of dysfunctional mitochondria to maintain cellular homeostasis^49^. However, it is also observed that excessive or dysregulated mitophagy can contribute to chondrocyte apoptosis following cartilage degradation ^26^. Our study showed that the p62-mediated mitophagy inducer had degenerative effects on goat TMJ cartilage cells, and this effect was not observed when cells were treated with LOXL2. Both these studies identified the unique role of LOXL2 in protecting TMJ cartilage by enhancing mitochondrial function. Overall, LOXL2 showed a protective effect on chondrocytes as well as mitochondria. Integrin-focal adhesion kinase (FAK) signaling is required for triggering the NF-κβ pathway ^50, 51^. However, one published study showed conflicting results that LOXL2 activated Integrin-FAK signaling in mandibular chondrocytes^52^. Our data also showed *Loxl2* knockout mice TMJ has an increase in Integrin signaling whereas LOXL2 overexpression reduced it.

The limitations of this study are that the findings are evaluated in murine and goat tissues, and whether LOXL2 protects TMJ cartilage during clinical studies by similar mechanisms is unknown.

In conclusion, *Loxl2* deletion promotes chondrocyte apoptosis by promoting IL-1β, NF-κβ, MMP13, and ADAMTS5, which are activated during synovitis, inflammation, and OA. However, Loxl2 deletion reduces the aggrecan, collagen, and proteoglycan network, which is critical in maintaining a healthy TMJ. LOXL2 overexpression could reverse these apoptotic changes, and pNF-κβ in murine and goat tissues point out its utility. The findings of this study have significant implications in developing new therapies for TMJ disorders. Further research will help us to fully understand the potential risks and benefits of using LOXL2 as a therapeutic target in TMJ-OA.

## Acknowledgment

This study was supported by an NIH grant R01 DE031413 (MB).

## Ethical Compliance

All procedures performed in this study involving human participants were performed according to the ethical standards of the institutional and/or national research committees.

## Author Contributions

Rajnikant Dilip Raut, Chumki Choudhury, Faiza Ali, Amit Kumar Chakraborty, Mohammed Moeenuddin Ahmed, Cheyleann Del Valle-Ponce De Leon, Harshal V. Modh, and Manish V. Bais performed the experiments and analyzed the data. Pushkar Mehra, Yuwei Fan, Alejandro Almarza, and Manish V. Bais helped with the conception, interpretation, and manuscript editing. Manish V. Bais conceived and designed the study. Rajnikant Dilip Raut and Manish V. Bais interpreted the data and wrote the manuscript.

## Data Availability

Raw RNA sequencing data were submitted to NCBI GEO under accession ID GSE276978 (mice) and GSE277139 (goat) (reviewer tokens will be available upon request). All other software and packages used in this study are described in the methods section.

## Supplementary Figure Legends

**Fig. S1: *Loxl2* knockout results in OA-like molecular changes.**

**a)** IPA *Loxl2* knockout genes normalized to control shows the predicted activation of cleavage of collagen fiber and proteoglycans, cartilage degradation and chondrocyte death. **b)** IPA showed the activation of IL-1β-related network genes. The upregulated genes within this network are hallmarks of cartilage degeneration and naturally progressive osteoarthritis.

**Fig. S2: IL-1β treatment results in OA-like molecular changes.**

**a)** DGE analysis confirms elevation of key OA genes, MMP13, ADAMTS5, and depletion of ACAN in both SLC and MLC with IL-1β treatment. **b)** IPA shows inhibition of cleavage of collagen fiber and proteoglycan, resulting in inhibition of cartilage degradation and activation of glycosaminoglycan (GAG) production. *p<0.05, **p<0.01, ***p<0.001, ****p<0.0001.

**Fig. S3: Protein and phosphoprotein expression analysis in vehicle, IL-1β, and IL-1β+LOXL2-treated goat TMJ chondrocytes.**

Contour plot representation of flow cytometry for MMP13, phosphorylated NF-κβ (pNF-κβ), and phosphorylated ERK1/2 (pERK1/2).

**Fig. S4: LOXL2 treatment in SLC identified its unique role in maintaining the mitochondria function:**

**a-b)** DESeq2 Log2 fold changes for common genes between *Loxl2* knockout and LOXL2 treated SLC in both MT and OP. Corresponding Wald test p-values are indicated on each bar. **c-d)** DESeq2 Log2 fold changes for common genes LOXL2 treated MLC in both MT and OP. Corresponding Wald test p-values are indicated on each bar.

**Fig. S5: LOXL2 treatment reduces IL-1β-induced chondrocyte apoptosis via inhibiting the NF-κβ pathway.**

**a)** IPA of *Loxl2* knockout RNA-seq data shows activation of IL-1 mediated NF-κβ activation leading to increased inflammation, cleavage of collagen fiber, and chondrocyte apoptosis. **b)** IPA of LOXL2 treatment from RNA-seq data shows inhibition of IL-1β mediated NF-κβ, leading to decreased inflammation, cleavage of collagen fiber, and chondrocyte apoptosis.

