## Supplementary figures and images for "LOXL2 Deletion Triggers TMJ Osteoarthritis While Overexpression Protects Against NF-κβ-Induced Chondrocyte Apoptosis"

### Supplementary figure 1

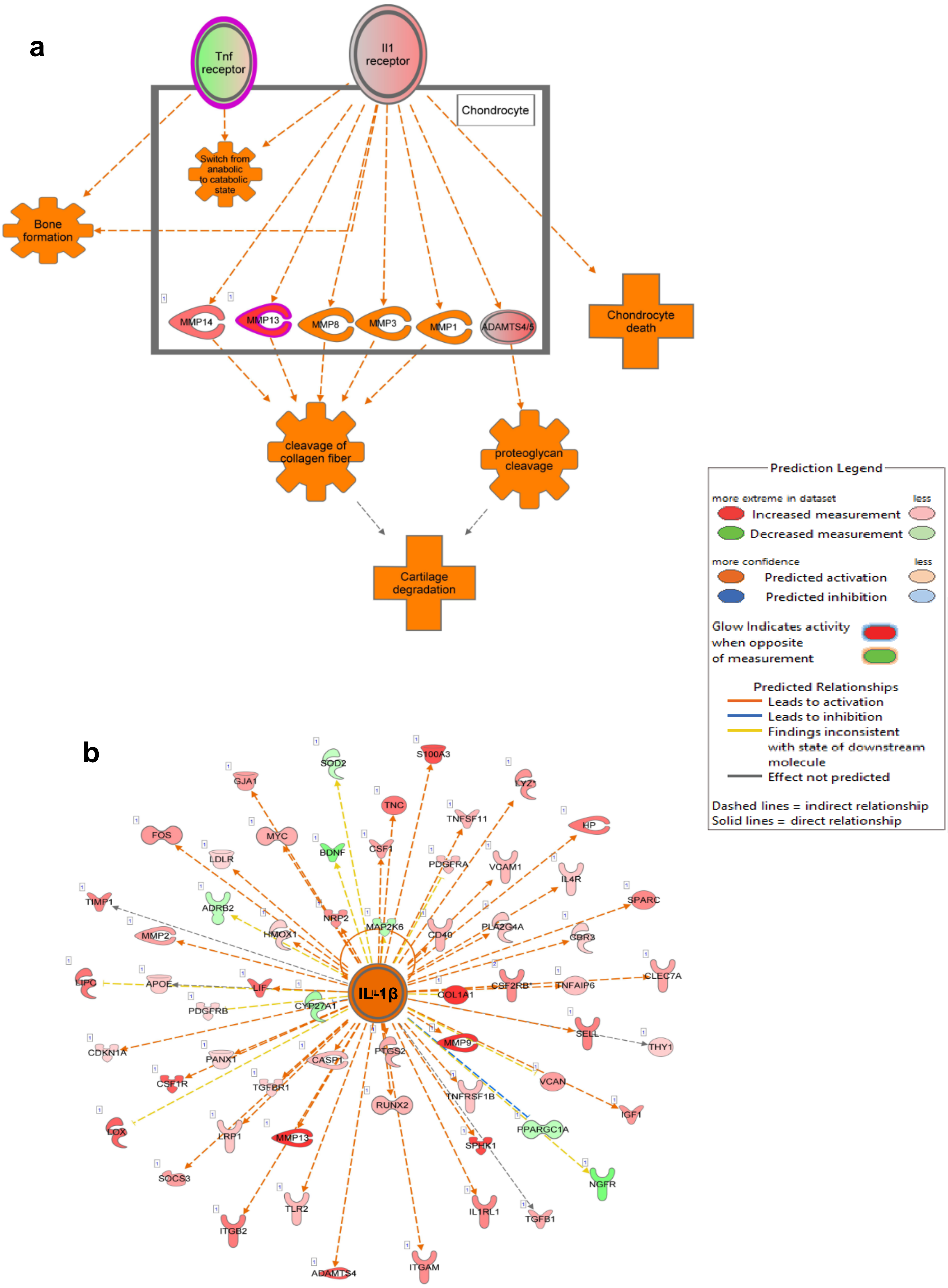

### Supplementary figure 3

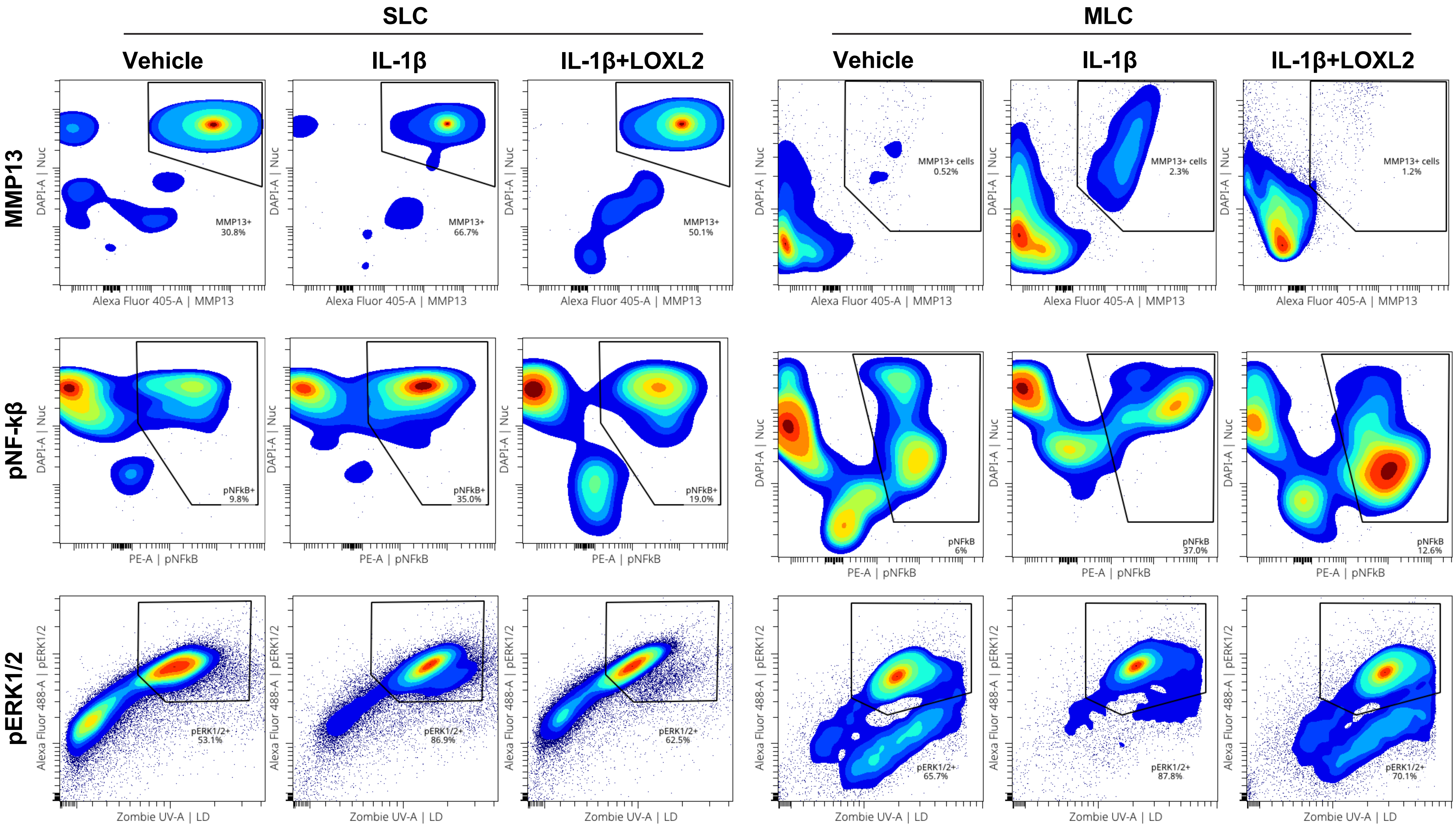

### Supplementary figure 4

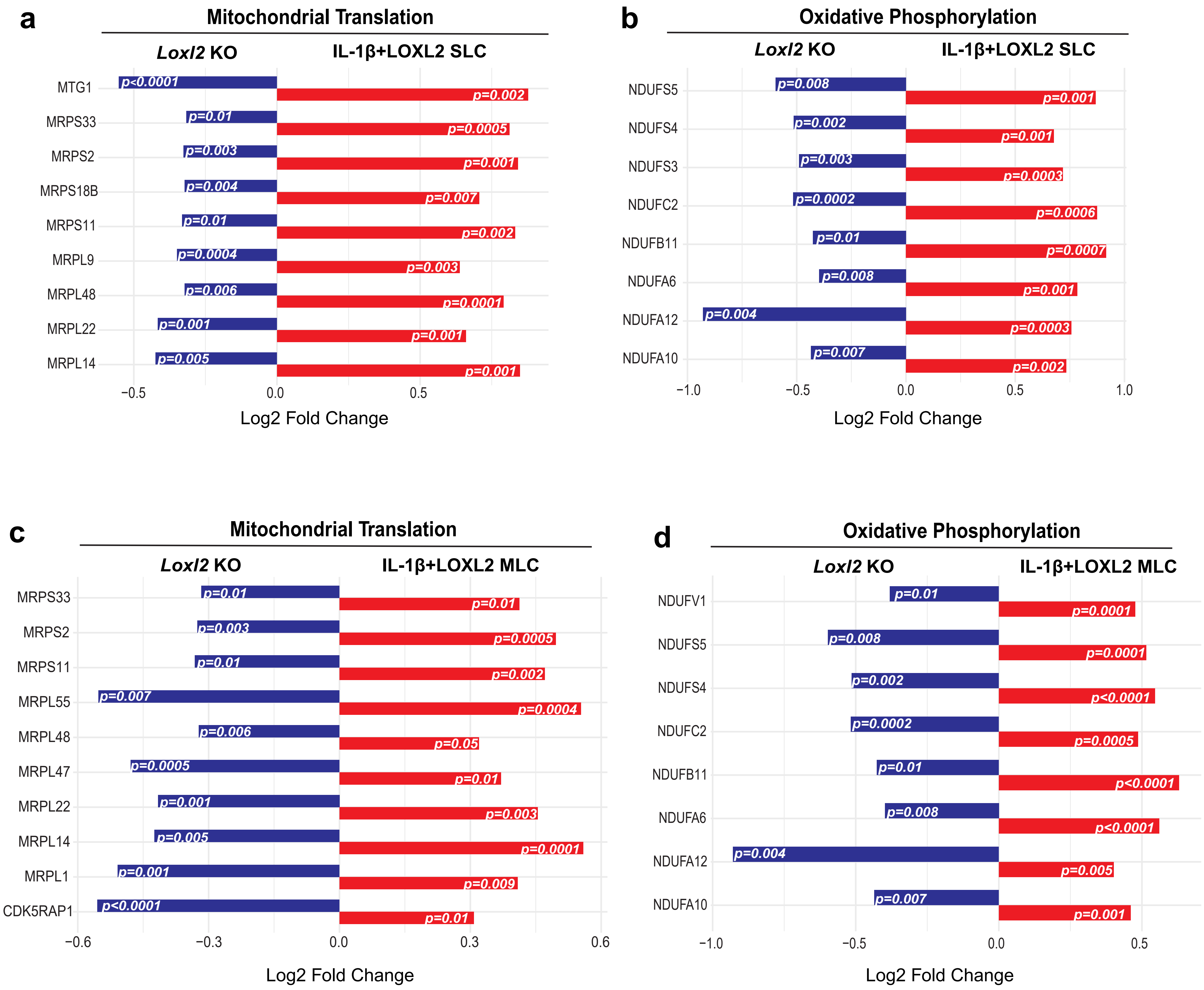

### Supplementary figure 5

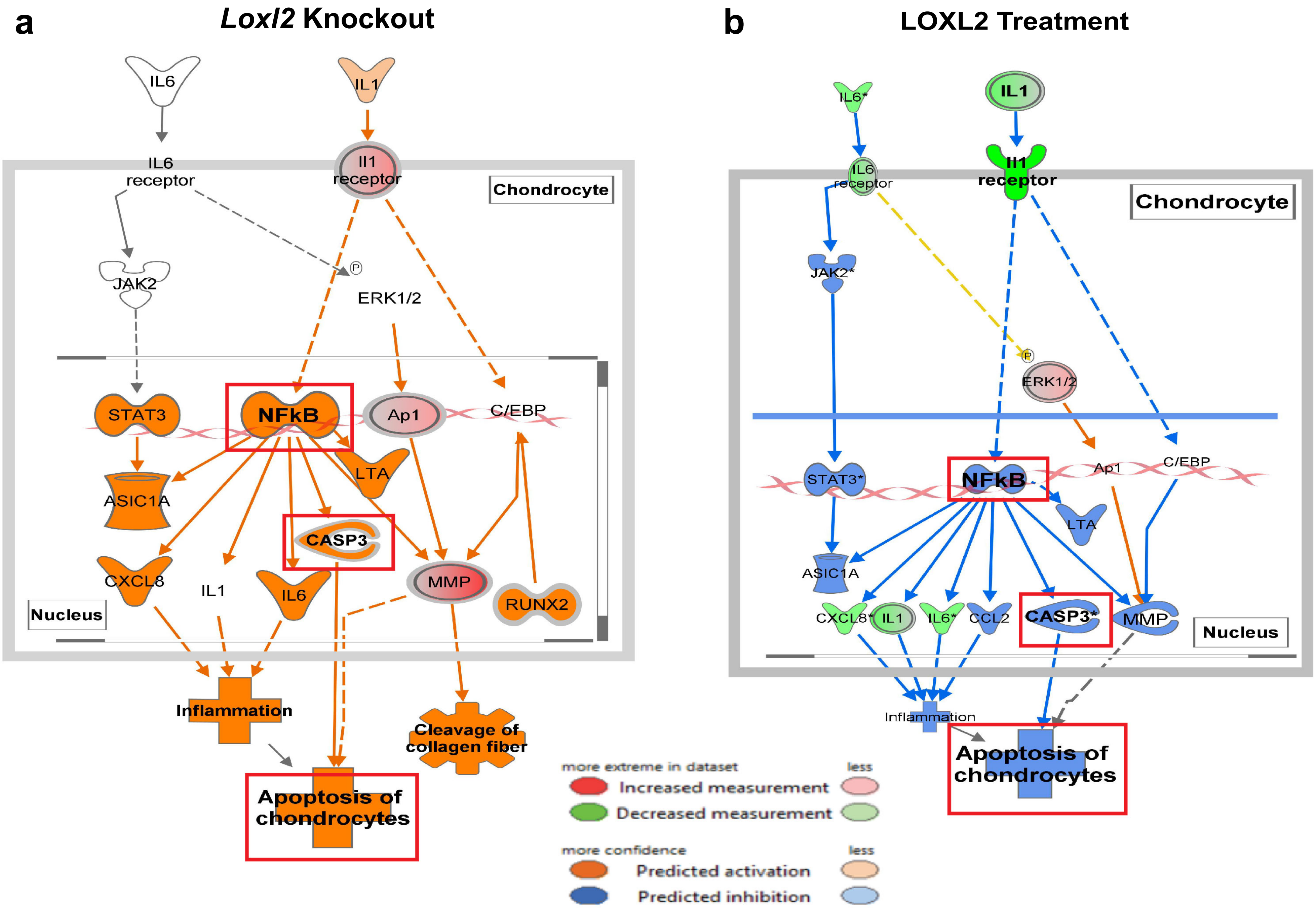
