## Supplementary figure 2 for "LOXL2 Deletion Triggers TMJ Osteoarthritis While Overexpression Protects Against NF-κβ-Induced Chondrocyte Apoptosis"

**Fig. S2**

**a** Expression of key OA genes upon IL-1β treatment against corresponding vehicle treatment

SLC

| gene_name | log2FC | pvalue | padj/FDR |
|---|---|---|---|
| MMP13 | 1.798689 | 0.000371 | 0.002732 |
| ADAMTS5 | 1.731681 | 1.55E-55 | 2.50E-53 |
| ACAN | -1.77845 | 3.89E-09 | 6.74E-08 |

MLC

| gene_name | log2FC | pvalue | padj/FDR |
|---|---|---|---|
| MMP13 | 7.439413 | 2.44E-16 | 9.73E-15 |
| ADAMTS5 | 2.492373 | 1.45E-17 | 6.38E-16 |
| ACAN | -1.27752 | 2.98E-29 | 2.70E-27 |

MLC: Middle cartilage layer-derived cells
SLC: Superficial layer-derived cells

**b**

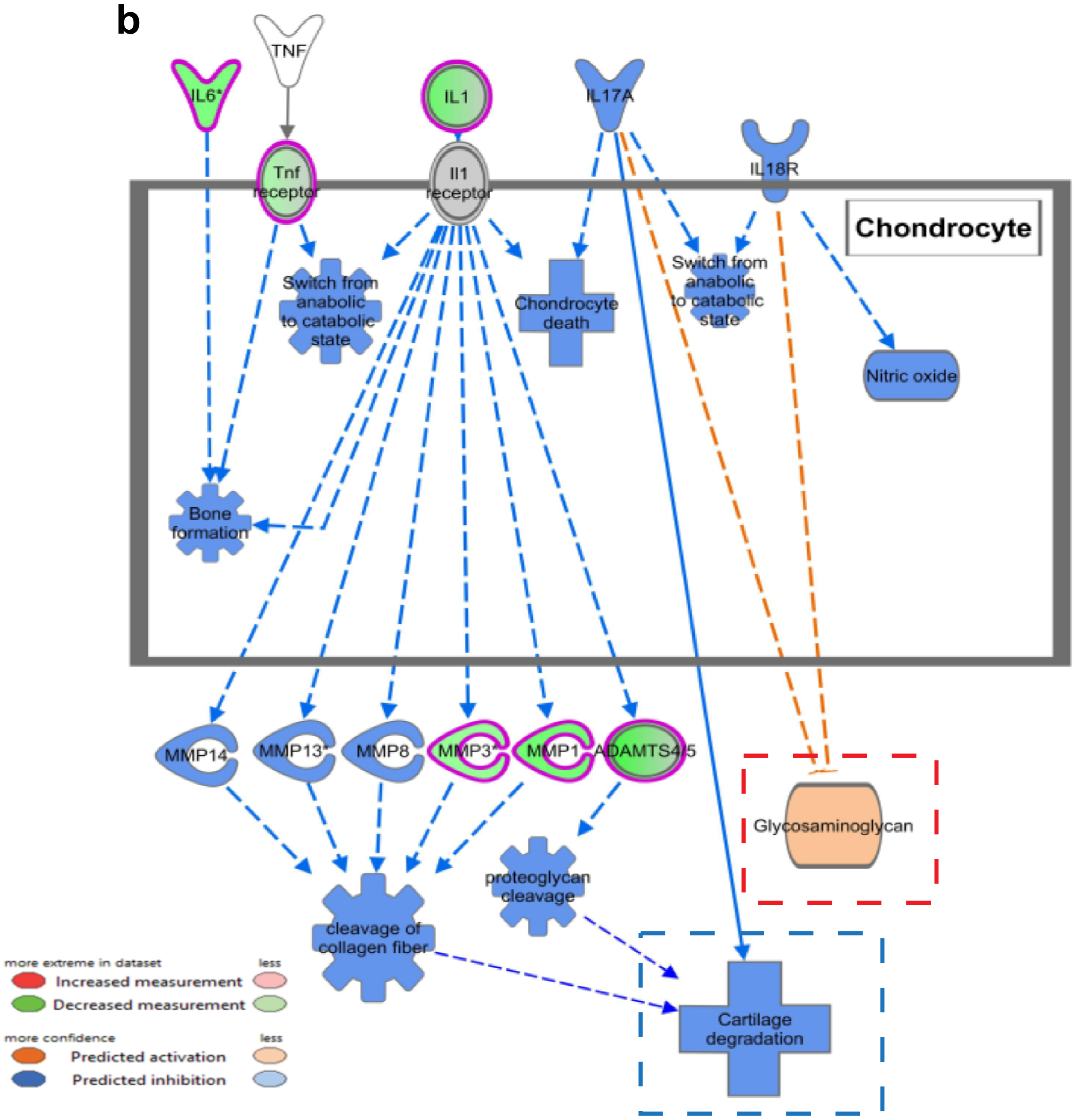
